# Automated Workflow for Instant Labeling and Real-Time Monitoring of Monoclonal Antibody N-Glycosylation

**DOI:** 10.1101/2022.12.22.521623

**Authors:** Aron Gyorgypal, Oscar Potter, Antash Chaturvedi, David N. Powers, Shishir P. S. Chundawat

## Abstract

With the transition toward continuous bioprocessing, process analytical technology (PAT) is becoming necessary for rapid and reliable in-process monitoring during biotherapeutics manufacturing. Bioprocess 4.0 is looking to build an end-to-end bioprocesses that includes PAT-enabled real-time process control. This is especially important for drug product quality attributes that can change during bioprocessing, such as protein N-glycosylation, a critical quality attribute for most monoclonal antibody (mAb) therapeutics. Glycosylation of mAbs is known to influence their efficacy as therapeutics and is regulated for a majority of mAb products on the market today. Currently, there is no method to truly measure N-glycosylation using on-line PAT, hence making it impractical to design upstream process control strategies. We recently described the N-GLYcanyzer: an integrated PAT unit that measures mAb N-glycosylation within 3 hours of automated sampling from a bioreactor. Here, we integrated Agilent’s Instant PC (IPC) based chemistry workflow into the N-GLYcanzyer PAT unit to allow for nearly 10x faster mAb glycoforms analysis. Our methodology is explained in detail to allow for replication of the PAT workflow as well as present a case study demonstrating use of this PAT to autonomously monitor a mammalian cell perfusion process at the bench-scale to gain increased knowledge of mAb glycosylation dynamics during continuous biomanufacturing of biologics using Chinese Hamster Ovary (CHO) cells.

**Graphical Abstract:** 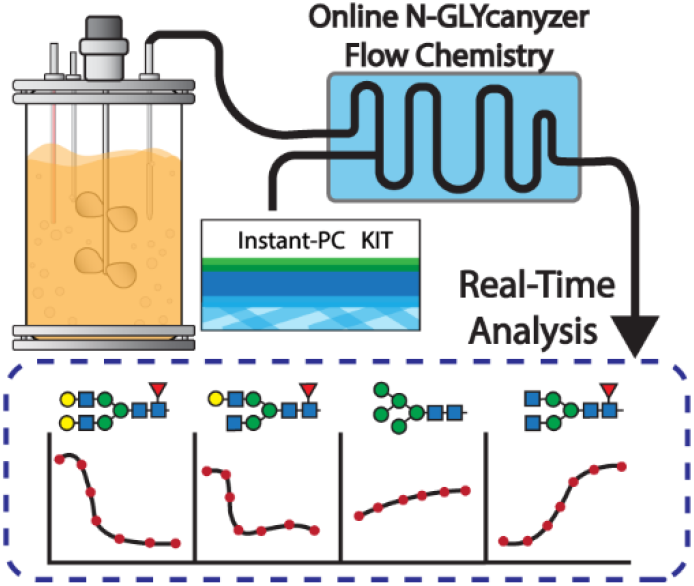

## Introduction

Implementation of advanced PAT and process control in the biopharmaceuticals and bioproducts manufacturing industries continues to lag behind the traditional petrochemical/chemical industry. The current goal towards bioprocess 4.0 is the creation of an end-to-end integrated bioprocess that runs, controls, and continuously improves the process following feed-back/forward control loops enabled by advances in automation and artificial intelligence^1,2^. However, due to inherent complexities in bioprocesses such as post-translational modifications of therapeutic proteins during biomanufacturing, the creation of PAT tools to continually monitor the critical quality attributes (CQA’s) of biologics is a challenge in itself^3–5^. A current bottleneck for both bioprocess and bioproduct characterization is the combination of high-throughput and autonomous PAT with high-resolution product quality analytics^6^.

N-linked glycosylation of proteins has garnered attention as a critical quality attribute for many biologic products, especially monoclonal antibodies (mAbs), as macro-heterogeneity in mAb glycoform structures are known to influence the pharmacokinetics, pharmacodynamics, and immunogenicity of the final drug product^7,8^. N-linked glycosylation is conserved for IgG monoclonal antibodies on their heavy chain at the Asn-297 site, with some products also having N-glycosylation in the variable region as well. The heterogeneity of N-linked glycosylation comes from the multitude of variations in the glycan branches due to the high number of sugar moieties possible as well as the specific linkages present that is influenced by the activity of different glycosidases and glycosyltransferases during cell growth, stationary, and death phases^8,9^. The glycosylation pattern tends to be also sensitive to the process parameters and the extracellular environment consequently. These parameters are known as critical process parameters (CPPs) and include cell culture temperature, pH, dissolved oxygen concentration, and agitation rate^10–14^. Because of this, a process must be well defined to make the glycosylation patterns reproducible between multiple batches^15^. Additional complexity is further added if the mAb product of interest is a biosimilar that has stricter tolerances for CQAs to match the originator or innovator drug product^16,17^.

Released glycan analysis often involves enzymatic deglycosylation of mAbs isolated from the cell culture using Peptide:N-glycosidase F (PNGase F), followed by glycan labeling by suitable fluorophore tag and labeled glycan enrichment using solid phase extraction (SPE). Traditional methods involve isolated mAb denaturation before a 4-24 hour incubation period for deglycosylation followed by an optional cleanup step to remove deglycosylated protein from the solution. Next, a 2-3 hour incubation step is necessary for fluorescently labeling the released glycan using reductive amination to conjugate a fluorophore like 2-aminobenzenamide (2-AB) to the reducing end of the glycans to increase analytical sensitivity. Finally, the excess label is then removed using solid phase extraction (SPE), and the sample is then dried and reconstituted into a suitable matrix before analysis by High-Performance Liquid Chromatography (HPLC) system coupled to a suitable Fluorescence Detector (FLD). This whole process can take anywhere from 2-3 days from start to end^18^. However, newer technology can allow this workflow to be further streamlined, such as using proprietary PNGase F kits to reduce deglycosylation reaction times to under few minutes, as well as using instant labeling chemistries that allow for nearly instantaneous fluorophore-glycan conjugation. Such technologies can condense the overall N-glycan release and sample prep workflow to less than one hour ^19^. Examples of such proprietary chemistry kits include Agilent’s AdvanceBio Gly-X Technology, as well as Waters’ GlycoWorks RapiFluor-MS^20,21^. While such recent innovations have been able to speed up sample preparation time as well as increase throughput using a 96-well plates design, these kits are not suitable for in-process real-time testing during manufacturing and are more suitable for quality control (QC) based analysis^22^.

Here, we look to enable rapid near real-time analysis of mAb N-glycans by integrating the Agilent Gly-X Instant Procainamide (IPC) chemistry and workflow into the N-GLYcanyzer PAT system. This will allow for faster mAb glycoforms analysis during bioprocessing compared to the traditional 2-AB labeling approach that was recently reported^23^. We also show the utility of using the IPC tag to deconvolute glycan peaks using at-line integrated liquid chromatography based mass spectrometry (LC-MS). We demonstrate how instant IPC chemistry can be integrated into an online PAT workflow for automated analysis of mAb glycoforms. Finally, we highlight a case study demonstrating the utility of this automated PAT workflow to rapidly monitor mAb glycoforms produced by a CHO cell perfusion bioprocess.

## Materials and Methods

### Cell line and shake flask cell culture

The Chinese Hamster Ovary (CHO-K1) cell line producing a recombinant trastuzumab, a biosimilar for Herceptin, was kindly donated by GenScript Biotech Corporation (Piscataway, NJ). A seed train was started by thawing one ampule of cells (10×10^6^ cell/mL) from the working seed bank into high intensity perfusion CHO (HIP-CHO) medium (Thermo Fischer Scientific, Waltham, MA) into a 125 mL unbaffled shake flask (VWR, Radnor, PA) with a 40 mL working volume to a seed density of 0.5×10^6^ cells/mL containing 0.1% anticlumping agent (Thermo Fischer Scientific, Waltham, MA). The cells were grown at 37°C, 130 RPM, and 8% CO_2_ in a New Brunswick S41i CO_2_ Incubator (New Brunswick Eppendorf, Hamburg, Germany) for 4 days and passaged twice to 0.5×10^6^ cell/mL into a 250 mL shake flask and then into a 500 mL shake flask, and then grown for 4 days before inoculation into the bioreactor.

### Perfusion bioreactor cell culture

The bioreactor cell culture experiments were conducted in a 3L glass bioreactor using Biostat B-DCU controller (Sartorius, Göttingen, Germany) with a working volume of 1.75L. Temperature and pH control was initiated before inoculation and set at 37°C and pH 7.1, respectively. Dissolved oxygen (DO) was also brought to a setpoint of 50% DO. The pH was controlled by sparging either CO_2_ or by bolus additions of 0.5M NaOH (Sigma Aldrich, St. Louis, MO). The bioreactor was inoculated to an initial density of 0.5×10^6^ cells/mL. Offline samples were taken daily to analyze various culture parameters (e.g., glucose, lactate, glutamate, glutamine, Na, K, Ca) on a BioProfile Flex2 Analyzer (Nova Biomedical, Waltham, MA). Product titer was analyzed offline from spent media daily by protein A chromatography on the Agilent Bioinert 1260 HPLC system using a Bio-Monolith Recombinant Protein A column (Agilent Technologies, Santa Clara, CA). An XCell^™^ ATF system (Repligen, Waltham, MA) was used for steady-state perfusion slowly ramping up the exchange rate from 0.25 to 1.0 vessel volumes a day (VVD) between day 4 and day 8. The bleed rate was also adjusted proportionally with the permeate rate using the pumps to maintain a constant VVD and cell viability throughout the culture duration.

### Off-line N-glycan sample preparation and analysis

Offline N-glycan analysis was done using AdvanceBio Gly-X N-glycan prep with InstantPC (GX96-IPC, Agilent Technologies, Santa Clara, CA) following the manufacturer’s instructions. Briefly, spent media was removed from the bioreactor daily and the sample was purified using a Protein A HP SpinTrap (Cytiva, Marlborough, MA) with 20mM phosphate buffer pH 7.2 as a binding buffer and 0.1% formic acid as the eluent. The sample was then neutralized using 1M HEPES Solution pH 8.0 to a neutral pH before buffer exchange into 50 mM HEPES solution pH 7.9 and then concentrated to ^~^2 g/L using a 10 kDa MWCO spin column (VWR, Radnor, PA). Next, 2 μL of Gly-X denaturant was added to 20 μL of the sample prior to heating it to 90°C for three minutes. After cooling, 2 μL of N-Glycanase working solution (1:1 Gly-X N-Glycanase, Gly-X Digest Buffer) was added, mixed, and incubated at 50°C for five minutes. Afterward, 5 μL of Instant PC Dye solution was added, mixed, and incubated for an additional 1 minute at 50°C. The sample was then diluted with 150 μL of load/wash solution (2.5% formic acid, 97.5% acetonitrile (ACN)). Next, 400 μL of load/wash solution was added to the Gly-X Cleanup Plate along with the ^~^ 172 μL of sample. A vacuum was applied (<5 inches Hg) until the sample passed through. Samples were then washed twice with 600 μL of Load/Wash solution before being eluted into a collection plate with 100 μL of Gly-X InstantPC Eluent with vacuum (<2 inches Hg). These samples were run on a 1260 Infinity II Bio-Inert LC System (Agilent Technologies, Santa Clara, CA) using an AdvanceBio Glycan Mapping column 2.1 × 150 mm 2.7 micron (Agilent Technologies, Santa Clara, CA). Mobile phase A was 50 mM ammonium formate adjusted to pH 4.4 using formic acid and mobile phase B was acetonitrile. The flow rate was set to 0.5 mL/min, and FLD was set to ex. 285 nm/ em. 345 nm, column temp was at 55°C. The initial eluent was held at 80% B for 2 minutes then dropped immediately to 75% B. From 2 minutes to 30 minutes the eluent was changed from 75% B down to 67% B in a linear gradient, and then from 30 to 31 minutes it was decreased from 67% B down to 40% B. From 31 to 33.5 minutes the ACN concentration was brought back to 80% at which level it was held until the end of the run at 45 minutes. Relative abundances of individual glycoforms was done on OpenLab CDS v3.5 (Agilent Technologies, Santa Clara, CA).

### Automated mAb titer analysis using N-GLYcanyzer

Titer was checked at least once a day using the N-GLYcanyzer system using the ProSIA subunit (FIAlab Instruments, Seattle, WA) following the method described in a previous study^23^. Briefly, bioreactor supernatant was pumped from the bioreactor through a filtration membrane and sent to the ProSIA system that integrated with a miniature protein A column. The column was machined in-house using PEEK (polyether ether ketone) material with an inner diameter of 2 mm and length of 30 mm and was packed with MabSelect SuRe Protein A resin (MilliporeSigma, Burlington, MA). Once mAb was adsorbed on the column the samples were washed with 20 mM phosphate buffer pH 7.2 and then eluted using 200 μL of 0.1% formic acid. The eluted sample was sent through an in-line UV spectrometer (Ocean Optics, Dunedin, FL) that was integrated downstream of the Protein A column, measuring at 280 nm wavelength. The integrated peak was used to calculate protein titer against a 7-point calibration curve (MedChemExpress, Monmouth Junction, NJ). If the concentration was found to be sufficient, the sample of purified mAb was then used for released glycan sample preparation (as described below). However, if the concentration was found to be too low for the optimized automated N-GLYcanyzer method analytical range (i.e., less than 100 μg mAb in eluent), a larger cell-free sample was automatically drawn from the reactor and purified to increase the amount of purified mAb. The sample with desired concentration was then sent to the second subunit (N-GLYprep) for further sample preparation of the glycans from mAb.

### Automated N-glycan preparation and analysis using N-GLYcanyzer

**Scheme 1** depicts the overall workflow (Scheme 1A) and the flow path of the N-GLYcanyzer system (Scheme 1B). After mAb protein purification, glycan analysis was initiated on the N-GLYprep subunit as shown in scheme 1B. The sample was eluted from the Protein A column (having a volume of 200 μL as described above) and neutralized with 20 μL of 1M HEPES solution, pH 8. The neutralized sample was then homogenized within the syringe pump and all but 40 μL was sent to waste. Homogenization was done by aspirating and dispensing the sample to and from the syringe pump through a clear waste line. The remaining 40 μL homogenized sample was mixed with 4 μL of Gly-X denaturant, dispensed to the 90°C heated coil for 3 minutes, then aspirated back to the syringe pump to allow it to cool to room temperature. For deglycosylation, 4 μL of a N-Glycanase working solution was aspirated to the sample in the syringe, homogenized, and dispensed to the 50°C heated coil for 5 minutes and then aspirated back into the syringe pump. Labeling was done by aspirating 10 μL of IPC label to the sample within the syringe and dispensing the sample to the 50°C heater for 1 minute and then aspirating back to the syringe. The sample was then homogenized, and all but 1 μL was dispensed to waste. The 1 μL sample was then diluted with 250 μL of the wash solution (80% acetonitrile, 20% water) by aspirating the wash solution into the syringe and allowing it to mix with the 1 μL of sample. The wash solution mixed sample was then loaded onto a 2.1 x 5 mm trapping column (821725-906, AdvanceBio Glycan Mapping Guard Column) placed on an external valve (G5631A, 1290 Infinity II Valve Drive, Agilent Technologies, Santa Clara, CA) and washed with another 250 μL wash solution before the external valve was switched in-line with the analytical HPLC column and the HPLC gradient was started.

**Scheme 1:**
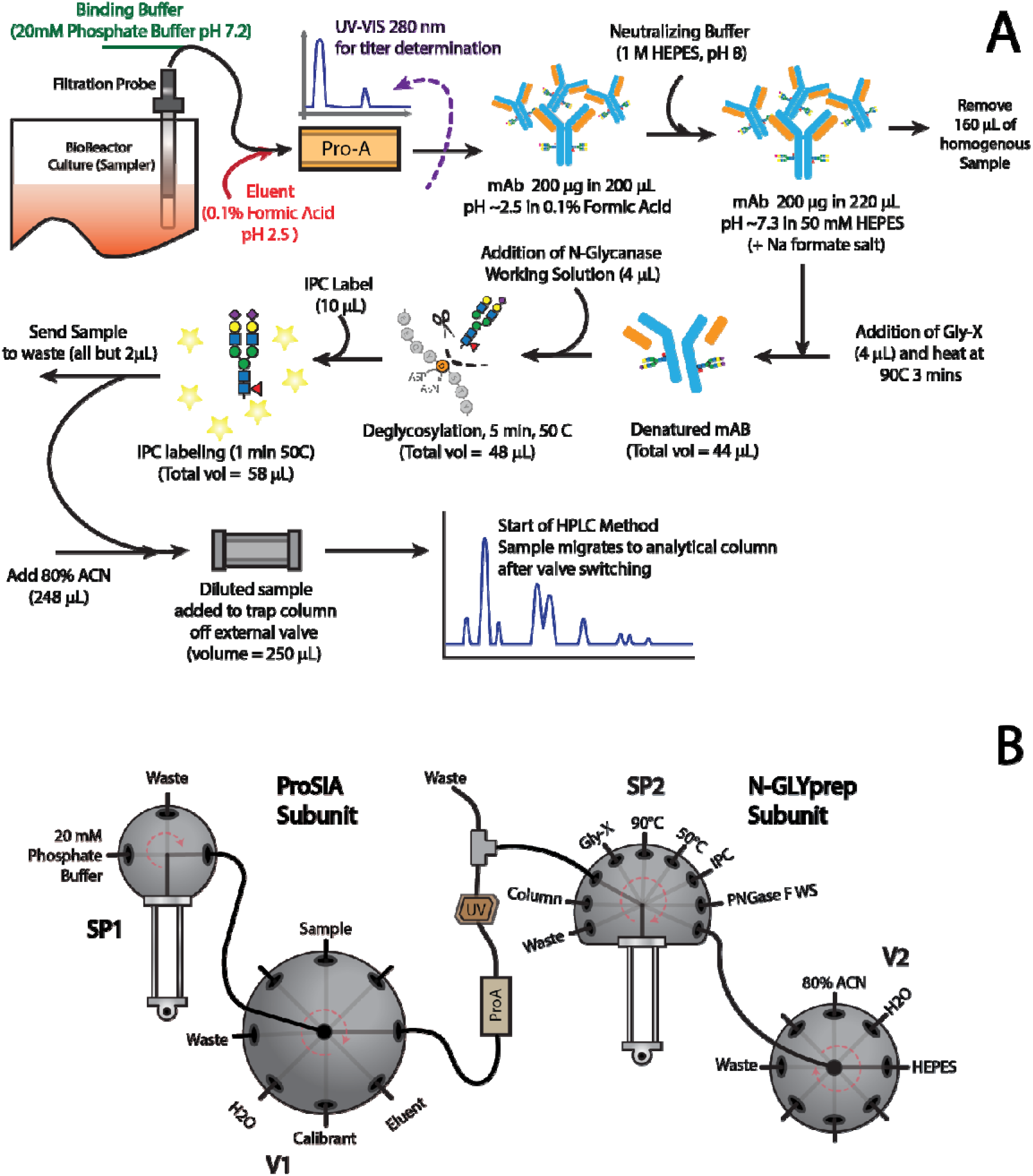
Instant PC glycan labeling chemistry workflow integration with N-GLYcanyzer PAT system. **(1A)** illustrates sample preparation process outlined within the methods section while **(1B)** shows the flow paths for sample preparation including syringe pumps 1 and 2 (SP1 and SP2, respectively) as well as the two associated valves (V1 and V2, respectively) within the overall workflow. The colors indicate different subunits: red indicates ProSIA system while blue indicates the N-GLYprep subunit, and gray is found in between the two subunits.

## Results and Discussion

### System Automation – Protein A Purification

The system used a 2 mm x 30 mm length column that was packed with MabSelect SuRe Protein A resin to purify the monoclonal antibody from the extracellular broth of the bioreactor culture. The binding buffer was 20 mM phosphate buffer pH 7.2 and the elution buffer was 0.1% formic acid. The column was conditioned before use. A fixed volume of cell-free reactor culture (200 μL) was removed using the filtration probe and pumped onto the protein A column. The sample was washed with the binding buffer before elution with 200 μL of 0.1% formic acid. During this time the eluent is monitored using UV 280 nm absorbance to calculate the mAb titer. If the concentration is too low for subsequent analysis the assay has been automated to be re-run at a higher sampling volume from the reactor to increase the final mAb concentration in the eluent. Afterwards the mAb eluent was neutralized to a pH of 7.9 – 8.0 using 20 μL of 1M solution of HEPES at pH 8.0. Prior experiments used a Tris-base solution for neutralization; however, it was found that tris-base interfered with the IPC labeling chemistry and was therefore discontinued for the online workflow. A sensitivity study was also run to measure the lowest limit of detection of the assay that are shown in **supplementary figure S1.** Based on the sensitivity study, we found that a mAb concentration as low as 0.1 g/L was sufficient for HPLC-FLD analysis, while 0.5 g/L gave better resolution of smaller eluting peaks. From this analysis it was decided that mAb would be concentrated to at least 0.5 g/L prior to glycan preparation post-protein A cleaning.

### System automation – deglycosylation and labeling

The integration of a bench-top assay based on manual steps into a flow-chemistry PAT system is non-trivial. Differences exist between the sample preparation for 2-AB and IPC based labeling chemistry. Labeling with 2-AB depends on a Schiff-base reductive amination of the released N-glycan reducing end moiety (after PNGase F treatment and spontaneous conversion of the glycosylamine product to a sugar aldehyde moiety) with the primary amine functional group of 2-AB forming an imine intermediate before reduction to a stable secondary amine. Conversely, the IPC method relies on a stable urea linkage formation between the instant procainamide label and the glycosylamine product formed immediately after PNGase F cleavage. This glycosylamine is unstable under non-alkaline conditions, losing its primary amine group which is necessary for the urea linkage formation^24,25^. The reaction schemes are summarized in **scheme 2** showing the PNGase F enzymatic reaction along with the subsequent IPC versus 2-AB based released N-glycan chemical reactions.

**Scheme 2:**
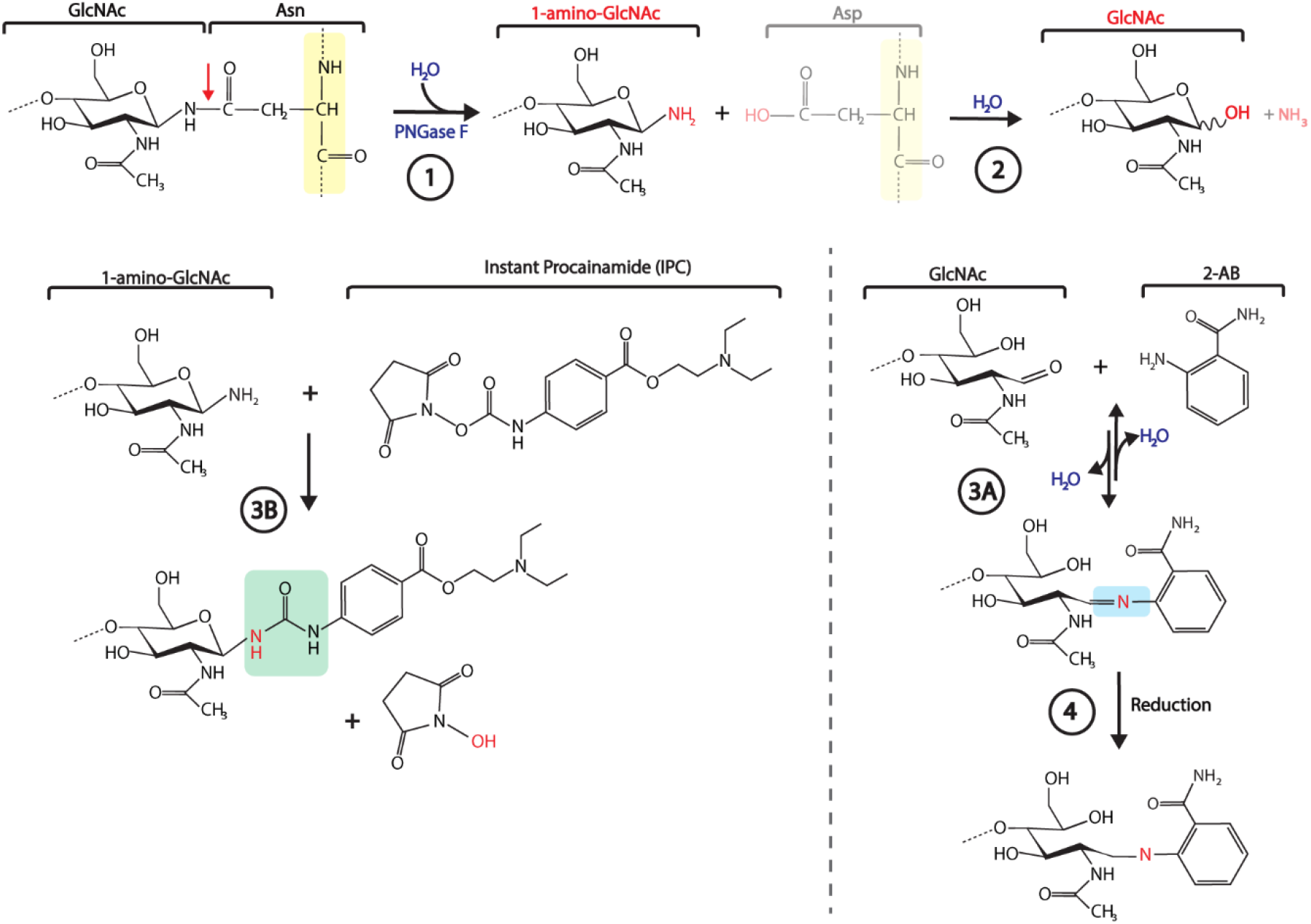
Reaction scheme associated with the enzymatic deglycosylation reaction followed by labeling chemistries using IPC versus 2-AB is shown here. In Step **(1)** the denatured antibody is treated with PNGase F that cleaves the innermost N-acetylglucosamine (GlcNac) of the N-glycan from the amino-acid backbone attached via the asparagine residue. This reaction releases the N-glycan oligosaccharide from the antibody protein backbone leaving a glycosylamine (1-amino-GlcNac) intermediate while converting the Asn residue to an aspartate (Asp) residue. The deglycosylated antibody is no longer needed for the subsequent reactions and is shown as faded in the reaction scheme. In Step **(2)** the reaction of the glycosylamine intermediate under slightly non-alkaline condition and prolonged reaction time in presence of water will lead to loss of an ammonia group that leaves behind the reducing sugar GlcNac intermediate. This free reducing sugar can be used as substrate for subsequent reductive amination reaction. In Step **(3A)** in the presence of a reactive amine such as 2-AB (a fluorophore) under high temperature and acidic reaction conditions the reducing sugar moiety of the N-glycan can react to form an imine intermediate (as shown highlighted in blue), which is unstable in water. In Step **(4)** the imine intermediate can be converted to a stable secondary amine in the presence of a strong reducing agent, and this final product is a N-glycan that is tagged with a 2-AB fluorophore. **(3B)** Conversely, the glycosylamine intermediate can instantaneously react with IPC to form a urea linkage (highlighted in green) under moderate reaction conditions leaving behind an N-Hydroxysuccinimide (NHS) by-product. Here the final product is a N-glycan that is tagged with a IPC fluorophore group.

A study was conducted to measure the labeling efficiency of IPC onto the glycosylamine as a function of PNGase F incubation time at two pH values: pH 7.5 and 8.0. The fluorescence intensity of G0F glycoform released from trastuzumab was monitored to examine the impact of PNGase F incubation time on relative concentration of glycosylamine intermediates release/labeled. This experiment provided some understanding of the relative amounts of glycosylamine intermediates formed after enzymatic cleavage to be readily available for IPC labeling. This provided insight to the optimum reaction time needed as PNGase F cleavage to release increasing concentration of glycosylamine intermediates was impacted by the subsequent hydrolysis of the intermediate to reducing sugars versus intermediate labeling by IPC probe. **Figure 1A** depicts representative chromatograms from the pH 7.5 assay condition. No bias was seen in the relative glycosylation pattern between all sample conditions and replicates for varying incubation times under either pH condition (data not shown). **Figure 1B** shows the integrated fluorescence intensity as arbitrary units (a.u.) of the most abundance labeled glycoform (G0F) as a function of the incubation time and pH. Interestingly, it was seen that in both cases the fluorescent intensity was high after 5 minutes of incubation, 12.94±0.67 a.u. at pH 7.5 and 12.65±0.32 a.u. at pH 8.0. The fluorescence intensity dropped at an incubation time of 10 minutes to 2.68±0.33 a.u. (pH 7.5) and 4.29±2.41 a.u. (pH 8.0). However, the fluorescence value was regained again under the pH 8 condition after 30 minutes to 14.41±1.42 a.u. and then stayed stable up to 30 minutes. However, under the pH 7.5 conditions, the fluorescence value stayed low even up to 30 minutes and then slowly increased to level off only after around 60 minutes to around 8.23 ± 1.53 a.u. While both conditions started with the same amount of substrate/enzyme (i.e., mAb and PNGase F), the amount of free glycosylamine available for the IPC reaction is almost twice as high at the higher pH reaction condition after one hour of incubation time. This can be attributed to the solution being slightly more alkaline and thus increasing free glycosylamine stability in solution prior to labeling with IPC.

**Figure 1.**
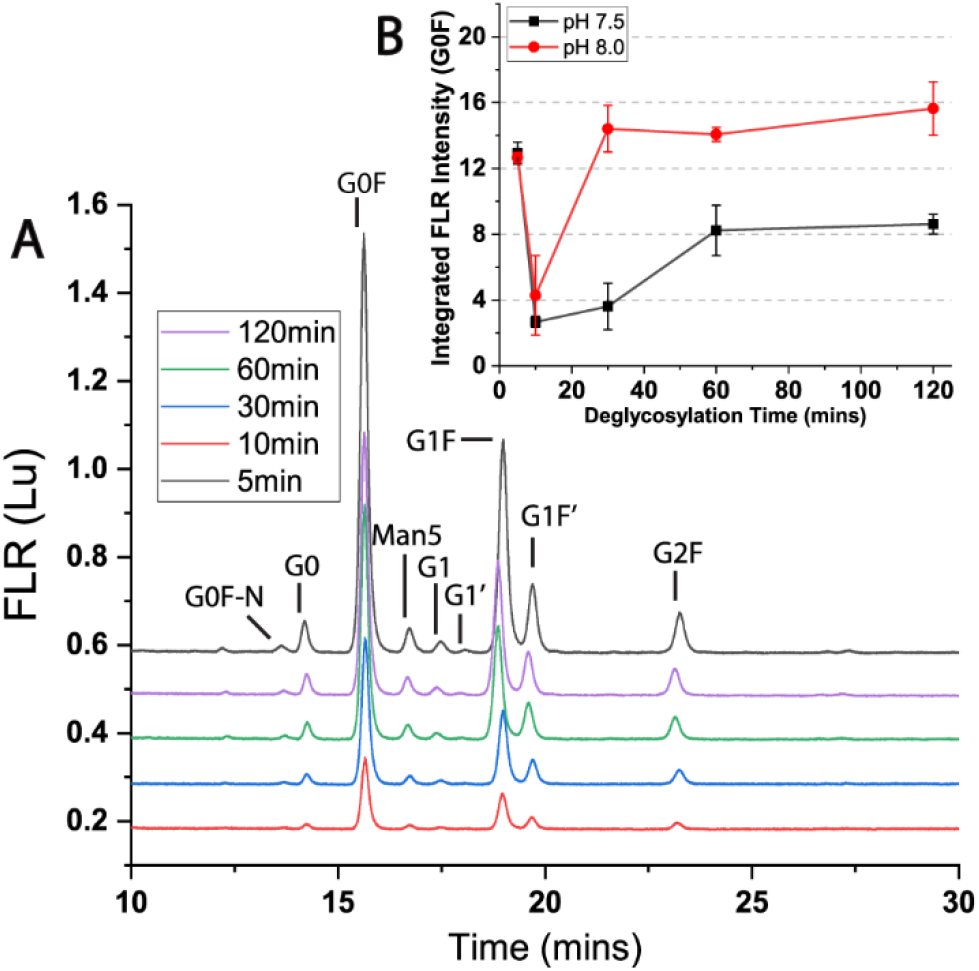
Impact on enzymatic deglycosylation reaction time on glycosylamine formation and labeling by IPC: Monoclonal antibody (^~^1 g/L) is buffer exchanged into HEPES solution at either pH 7.5 or pH 8.0 and then deglycosylated with PNGase F for varying incubation times from 5 minutes to upwards of 120 minutes (2 hours). Deglycosylated mAbs are all subjected to labeling with IPC immediately after their incubation times, cleaned offline, and then analyzed by HPLC-FLD. **(1A)** Representative chromatograms for the pH 7.5 reaction conditions showing changes in fluorescent intensity over reaction time is shown. **(1B)** Integration of the G0F glycoform to show changes in integrated fluorescence intensity between the two pH conditions over time showing an increase in the amount of labeled glycosylamine at the higher pH condition. All samples were run in n≤3 replicates.

To the best of our knowledge, there is no open literature that explains the dramatic decrease in free glycosylamine available to dye conjugation between the 5- and 10-minute reaction times. Furthermore, there are many potential unknowns in attempting to explain the mechanism behind this dynamic multi-step reaction kinetics behavior. For example, we still have limited knowledge of; (i) the extent of mAb denaturation that impacts subsequent PNGase F accessibility for glycan cleavage, (ii) the activity of PNGase F under varying pH conditions in the presence of the denaturant, and (iii) enzyme activity over time post initial burst phase as substrate available become rate-limiting. Earlier literature has characterized the kinetics of PNGase F, but not in the context of the glycosylamine formation and its subsequent degradation due to IPC labeling^26,27^. An additional unknown is the relative degradation rate of the intermediate glycosylamine to free-reducing sugar. Interestingly, there may be alternative chair confirmations of the glycosylamine that may be labeled as well^20^ as shown by Kimzey et al. within their application notes when first reporting on the IPC reagent for glycan labeling. Lastly, it is worth noting that pH does have an effect to the amount of glycosylamine available for IPC labeling, as it is known that the stability of glycosylamines is also pH dependent. While these arguments could explain the dynamic change in IPC labeled glycosylamine intermediate concentrations profile, further exploration was outside of the scope of the current project. In conclusion, to support automation and assay throughput we decided to use the 5-minute total incubation time for the enzymatic deglycosylation and IPC labeling step.

### HILIC trap column sample enrichment and injection

After glycans are deglycosylated and labeled with IPC, the samples must be purified to remove any excess label and other contaminants that may be present in solution. The offline, bench-top method uses a proprietary HILIC based material to remove such contaminants. This is done by diluting the labeled glycan samples with 0.1% formic acid in acetonitrile and then passing it through the proprietary HILIC material under vacuum, followed by three wash steps before eluting the bound glycan using a propriety eluent. For an online sample preparation methodology, the exact same steps cannot be easily replicated.

This problem was solved by instead introducing a small HILIC guard column to function as a trap column on a 6-port external valve off the HPLC, which acts as an extension to the analytical column upstream. This column functions as an enrichment step after IPC labeling and removes most contaminants without significant loss of all labeled glycans. Most of the labeled sample is sent to waste except for 1 μL which is diluted 1:250 with 80% acetonitrile and then injected into the trap column. Discarding the bulk of the labeled glycan sample facilitates adjusting the remaining solution to a weak HILIC eluent by addition of the 80% acetonitrile to better adsorb onto the HILIC trapping column. The remaining sample fraction is adequate because the fluorescence sensitivity of IPC labeled glycans is very high. The trapping column is then washed with another 250 μL of 80% acetonitrile. This six-port valve configuration can be seen in **Figure 2B.** For valve position 1 → 6: ports 1 and 4 contain the trapping column with port 5 as the inlet from the N-GLYcanzyer system allowing for the sample and wash solution to pass through to waste on port 6. In this valving position, the HPLC bypasses the trap column through ports 3 and 2. Once the sample is injected into the trap column and washed, the internal setting is switched to position 1 → 2 in which the trap column is now in-line with the HPLC mobile phase and the analytical column. At this point, the glycans on the trap column act as the extension of the analytical column and with the start of the mobile phase, the gradient decreases the concentration of the organic phase allowing for chromatography to take place. Surprisingly there was no peak broadening or peak shifting taking place with this online set-up and the chromatography for the online prepared samples ran nearly identically to the offline method prepared samples.

**Figure 2.**
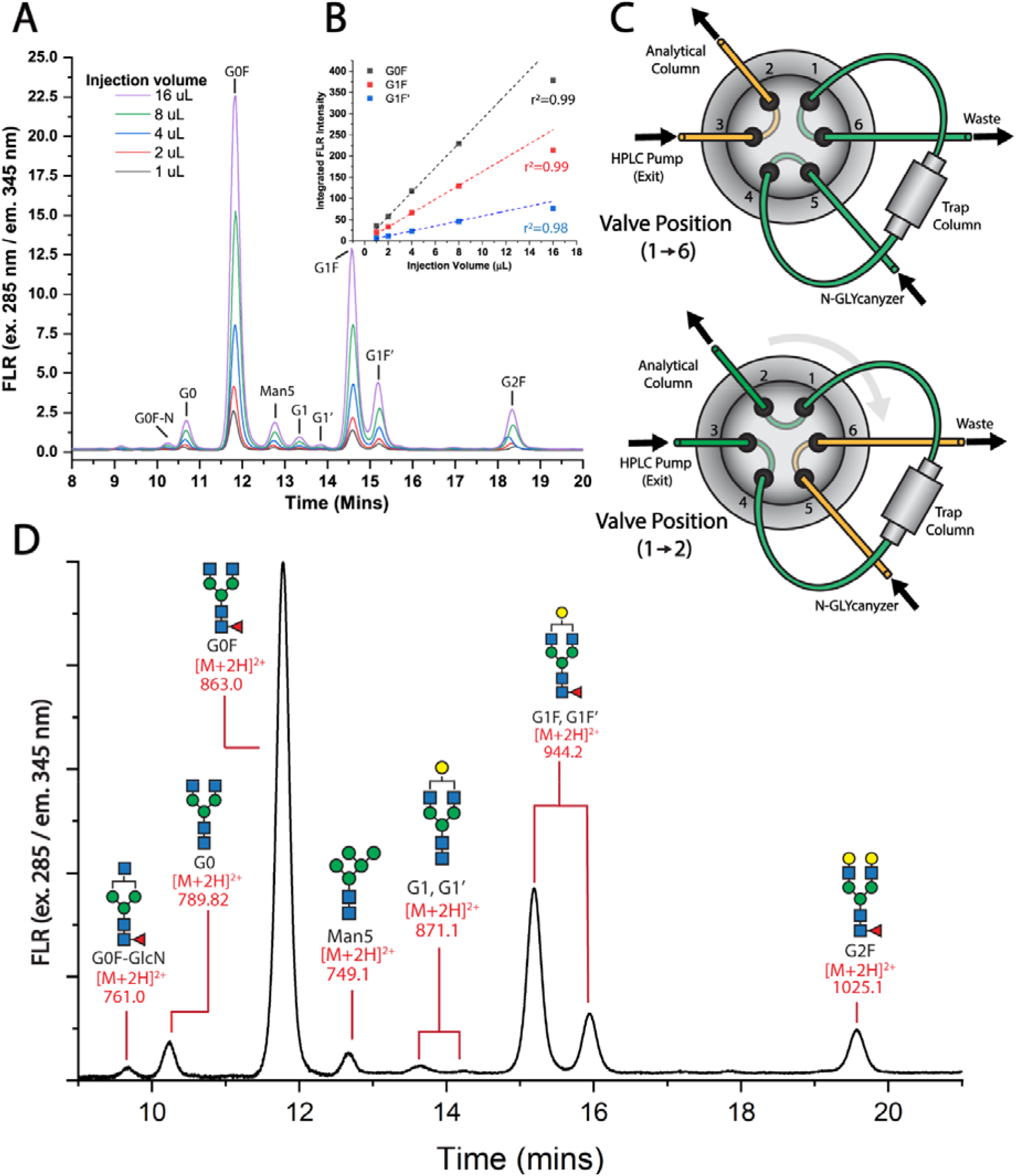
IPC labeled glycan sample cleanup using trap enables efficient labeled glycan separation on analytical column and detection using fluorescence and mass spectrometric detection methods: Increasing injection volumes of IPC labeled N-glycan sample does not cause bias towards relative trastuzumab glycoform abundances at lower injection volumes (ideally < 16 μl) onto the trap column. (2A) shows the injection and washing of different volumes of samples within a 250 uL matrix containing 80% acetonitrile and 20% water with no significant variation in residence time on column. **(2B)** Injection volumes for the three major glycoforms from the trastuzumab biosimilar, while Table 1 shows all glycoforms in tabulated form. **(2C)** The internal movement of external valve from “sample loading” valve position (1-6) to “HPLC analysis” valve position (1-2). The green lines represent the flow path taken by the sample during specific preparation and analysis steps. **(2D)** Example HPLC-FLD chromatogram of eluting glycoforms that were also confirmed using an offline LC-MS to indicate the specific mono-isotonic masses detected for each eluting glycan peak.

Next, we investigated the impact of sample injection volume onto the trapping column to understand the trapping efficiency or sample recovery. This was done by varying volumes of labeled glycan samples and diluting them to 250 μL before injection on to the N-GLYcanzyer unit. A sample of mAb around ^~^1 g/L was used for this experiment. The same sample was used for each injection to minimize batch-to-batch variability. Adjusting the injection volume and wash volume was also done to optimize this step, with 250 μL found to give the best cleaning efficiency versus glycan recovery (data not shown). **Figure 2B** shows the increase in fluorescence signal with the increase in prepared sample mass and is quantified for three of the most abundant glycoforms in **Figure 2C.** A linear response can be seen with the increase in sample mass up to 16 μL of loaded sample (r=0.99^2^), with linearity lost after 16 μL. This is quantified in terms of integrated fluorescence values as well as relative abundances in **Table 1A** and **1B.** There was no bias seen in the trastuzumab glycoform patterns upto 16 μL equivalent mass of sample injected onto the column. At the highest sample loading, there was a slight loss in linearity and the glycan distribution showed a decrease in relative abundances for the smaller glycoforms and a proportional increase for the larger glycoforms. For example, the relative abundance of G0F fell from 48.8% ± 0.1% to 42.6 % ± 0.2%, and G1F and G1F’ went from 27.5 ± 0.1% and 9.9% ± 0.0%, respectively to 32.1% ± 0.3% and 11.7% ± 0.2% relative abundances, respectively. This loss in retention and increase in recovery bias was expected for the highest loadings of samples tested. While increasing the sample loading volume (volume of sample in mostly aqueous buffer) the proportionality of the organic phase (acetonitrile concentration) will decrease leading to weaker retention of smaller glycoforms. Subsequently, larger glycans will tend to have stronger adsorption to the stationary phase causing a bias in sample recovery. Due to these results, we suspect that the trapping column was not overloaded at even the higher injection volumes/masses, but instead it is more likely that the weaker mobile phase caused bias in glycan retention^28,29^.

**Table 1.** Relative abundance of IPC-labeled trastuzumab glycoforms during sample recovery from trap column cleanup prior to analytical column injection. **(1A)** Absolute integrated peak fluorescent intensity, and **(1B)** relative absolute abundances of glycoforms from trastuzumab biosimilar at different injection volumes diluted into 250 μL 80% acetonitrile prepared for injection on trap column. All reported mean values are calculated with at least 2 technical replicates (n≤2). The standard deviations are also shown here.

| 1A | Glycan | Injection size (μL) (AVG+STD) n=2 |  |  |  |  |  |
| --- | --- | --- | --- | --- | --- | --- | --- |
|  |  | 1 | 2 | 4 | 8 | 16 | 32 |
| Integrated FLR Value | G0F-GN | 0.54 ± 0.02 | 0.95 ± 0.04 | 1.22 ± 0.01 | 2.41 ± 0.02 | 4.79 ± 0.02 | 5.76 ± 0.50 |
|  | G0 | 2.84 ± 0.05 | 4.62 ± 0.06 | 8.39 ± 0.03 | 15.96 ± 0.08 | 28.01 ± 1.27 | 32.57 ± 0.74 |
|  | G0F | 35.57 ± 0.16 | 57.16 ± 0.06 | 117.53 ± 0.67 | 229.92 ± 0.15 | 378.44 ± 3.63 | 484.24 ± 2.26 |
|  | Man5 | 2.24 ± 0.06 | 3.53 ± 0.14 | 7.22 ± 0.10 | 14.44 ± 0.25 | 23.10 ± 0.28 | 29.83 ± 0.26 |
|  | G1 | 0.91 ± 0.03 | 1.34 ± 0.06 | 2.85 ± 0.08 | 5.47 ± 0.20 | 8.39 ± 0.07 | 11.96 ± 0.13 |
|  | G1F | 20.50 ± 0.06 | 33.01 ± 0.13 | 66.94 ± 0.16 | 129.97 ± 0.12 | 213.37 ± 0.98 | 364.11 ± 7.14 |
|  | G1F' | 7.51 ± 0.03 | 11.53 ± 0.13 | 22.94 ± 0.11 | 45.98 ± 0.11 | 76.75 ± 0.48 | 132.56 ± 0.83 |
|  | G2F | 4.21 ± 0.03 | 6.46 ± 0.09 | 13.29 ± 0.04 | 26.77 ± 0.33 | 42.77 ± 0.40 | 75.00 ± 1.50 |

**Table 1. Relative abundance of IPC-labeled trastuzumab glycoforms during sample recovery from trap column cleanup prior to analytical column injection.**
| 1B | Glycan | Injection size (μL) (AVG+STD) n=2 |  |  |  |  |  |
| --- | --- | --- | --- | --- | --- | --- | --- |
|  |  | 1 | 2 | 4 | 8 | 16 | 32 |
| Relative Abundance | G0F-GN | 0.7% ± 0.0% | 0.8% ± 0.0% | 0.5% ± 0.0% | 0.5% ± 0.0% | 0.6% ± 0.0% | 0.5% ± 0.0% |
|  | G0 | 3.8% ± 0.0% | 3.9% ± 0.1% | 3.5% ± 0.0% | 3.4% ± 0.0% | 3.6% ± 0.1% | 2.9% ± 0.0% |
|  | G0F | 47.9% ± 0.0% | 48.2% ± 0.1% | 48.9% ± 0.1% | 48.8% ± 0.1% | 48.8% ± 0.1% | 42.6% ± 0.2% |
|  | Man5 | 3.0% ± 0.1% | 3.0% ± 0.1% | 3.0% ± 0.1% | 3.1% ± 0.0% | 3.0% ± 0.1% | 2.6% ± 0.0% |
|  | G1 | 1.2% ± 0.0% | 1.1% ± 0.1% | 1.2% ± 0.0% | 1.2% ± 0.0% | 1.1% ± 0.0% | 1.1% ± 0.0% |
|  | G1F | 27.6% ± 0.0% | 27.8% ± 0.1% | 27.8% ± 0.0% | 27.6% ± 0.0% | 27.5% ± 0.1% | 32.1% ± 0.3% |
|  | G1F' | 10.1% ± 0.1% | 9.7% ± 0.1% | 9.5% ± 0.0% | 9.8% ± 0.0% | 9.9% ± 0.0% | 11.7% ± 0.2% |
|  | G2F | 5.7% ± 0.1% | 5.4% ± 0.1% | 5.5% ± 0.0% | 5.7% ± 0.1% | 5.5% ± 0.0% | 6.6% ± 0.1% |

**Figure 3D** show the monoisotopic masses for each trastuzumab glycoform tagged with IPC and analyzed by LC-MS. While IPC is a fluorophore it also contains a tertiary amine which facilities IPC labeled species ionization in positive mode electrospray ionization mass spectrometry (ESI-MS). The utility of the IPC tag for MS analysis is showcased here to facilitate the concept of using this system with an LC-MS to allow for unknown labeled glycan mass identification. Samples of trastuzumab were analyzed on an offline LC-MS using a slightly longer gradient to allow for increased chromatographic separation prior to MS detection. The LC system was identical as before while the MS system was an Agilent Ultivo Triple Quadrupole mass spectrometer that is compatible with the overall N-GLYcanyzer workflow.

**Figure 3.**
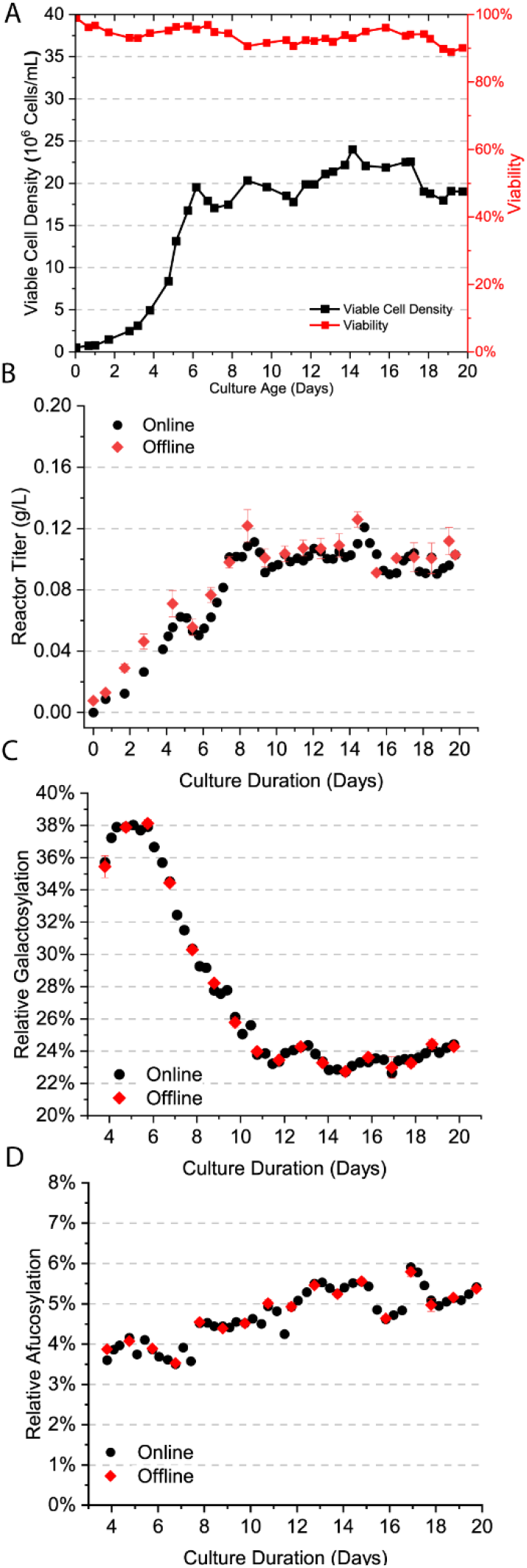
Online (in black) versus offline (in red) analysis of continuous perfusion bioreactor for mAb titer and major glycan Indices are shown here. **(3A)** Viable cell density and viability over cell culture. **(3B)** Reactor mAb titer, **(3C)** Relative mAb galactosylation, and **(3D)** Relative mAb afucosylation for trastuzumab. Here, online analysis was done using the integrated N-GLYcanyzer PAT system employing the IPC workflow, while offline analysis was done using standard offline methods.

A similar workflow was proposed by Bénet et al. as an online methodology to clean 2-AB using a trap column^30^. This workflow injected an impure 2-AB labeled glycan reaction mixture onto an HPLC and trapped the glycans on a BEH amide packed trap column using a 75% acetonitrile isostatic flow for a fixed amount of time to wash the trap column of contaminants while retaining the oligosaccharides before changing the valve position in line with the analytical column. This valve position change reversed the flow on the trap column as it eluted onto the analytical column. However, in our design, we did not change the flow on the trap column. The previous online clean-up workflow was comparable with offline cleaning to remove excess 2-AB as well. Our work shows a similar approach can be adopted using IPC tag over the 2-AB tag while using a superficially porous HILIC trap column.

Ultimately, it was found that a large volume of glycan sample can be loaded onto the trapping column without causing bias during downstream analytical chromatography step for LC-FLD or LC-MS. The assay was optimized so that around 2 μL of prepared labeled glycan sample will need to be diluted to 250 μL for the final online assay. This volume was chosen based on an analytical sensitivity criteria in case there is any loss in syringe aspiration and dispension tolerances used during the bioprocess campaign. An increase in the sample volume (pre-dilution) would need only be considered if fluorescent response was found to be low, depending on the mAb glycoform relative composition.

### Perfusion-based cell culture mAb glycoforms analysis

To showcase the utility of the N-GLYcanyzer system integrated with the IPC chemistry workflow, we studied a perfusion bioprocess producing a trastuzumab biosimilar. Perfusion mode of operation can become challenging to measure glycoforms since the mAb titers are considerably lower than that of a fed-batch counterpart as the product is constantly being harvested and cells are being bled to maintain a pseudo-steady state. Titer was measured every day starting at day 0, with glycoform analysis only started once a detectable concentration of mAb was seen in the culture on day 4. Culture harvesting was also started on day 4 at a 0.25 VVD, and cell bleeding started around day 6 as the viable cell density approached 20 million cells/mL. At this point, the perfusion rate was changed to 1.0 VVD with the bleed, and harvests were changed proportionally to maintain a semi-constant cell density throughout the 20-day culture.

**Figure 3A** shows the viable cell density and viability over the 20-day cell culture period. The viability stayed above 90% throughout the culture run and viable cell density maintained roughly between 18 and 23 million cells/mL. **Figure 3B** shows the titer monitored within the reactor throughout the culture using the online N-GLYcanyzer system as well as the standard offline analysis method. The measurements were taken once a day until day 4 and then roughly every 8 hours using the N-GLYcanyzer system. The offline measurements were done by taking at least two technical replicates (n=2), and the online measurements were done once per analysis. The titers measured under both systems showed very similar trends, with the offline measurements giving a slightly higher concentration. Once at a steady state the mAb space-time yield (STY) remained between 0.08 and 0.12 g/L/day through the culture.

The glycan indices (GI) calculated based on all detected trastuzumab glycoforms are shown in **Figure 3C** and **Figure 3D.** The relative galactosylation index was measured and calculated by the summation of all galactosylated glycoforms divided by the summation of all glycoforms, giving the relative level of mAb glycoforms galactosylation within the reactor. The results follow a similar trend as seen in our recent publication^23^ where the galactosylation rate tends to be high through the first few days of production (i.e., ^~^38% rel. galactosylation) and then sharply declines once the cells reach a pseudo-stationary phase (e.g., ^~^24% rel. galactosylation). The relative galactosylation rate is still within the quality tolerances set by the FDA based on a public release filing for a trastuzumab biosimilar^31^. The relative afucosylation index increased over time from around 4% to 5.5% at the end of the culture. This afucosylation index would be technically out of specification for a US-trastuzumab biosimilar based on a filing for another trastuzumab biosimilar as referenced above.

Madabhushi et al. proposed that the declining levels of relative mAb galactosylation are caused by an increase in the cellular productivity of mAb that results in decreased residence time within the Golgi apparatus and hence incomplete addition of terminal sugars like galactose to the N-glycan backbone^32^. Using the N-GLYcanzyer PAT system, it will be possible in the future to understand the dynamic changes in mAb N-glycosylation more frequently to better quantify the rates of change over time, and develop novel process control strategies to achieve bespoke mAb glycoform profiles. Incorporating a similar PAT system with an advanced multi-omics approach can also help reveal subtle changes within cellular pathways to gain a fundamental understanding of the metabolic bottlenecks impacting protein glycosylation.

## Conclusion

In this study, we have integrated a commercially available N-glycan release/labeling kit chemistry into the N-GLYcanzyer flow chemistry PAT system. This proof of concept system allows for real-time monitoring of a bioprocess to monitor protein N-glycosylation which could allow for future implementation of advanced control strategies during industrial-scale biologic biomanufacturing. The chemistry of glycosylamine formation during the enzymatic deglycosylation step was studied to understand how the relative formation and degradation rates over time at varying pH’s impact analytical sensitivity. A trapping column was introduced to the PAT flow system to allow for more accurate IPC labeled glycan capture, enrichment, and injection into a U/HPLC analytical column for fluorescence or mass spectrometric based product detection. The trap column was also characterized by exploring the sample matrix impact on glycan trapping. Lastly, we used the N-GLYcanyzer for automated real-time glycan analysis during a bench scale perfusion bioprocess to demonstrate the utility of the PAT system to measure changes in mAb glycosylation over time, especially monitoring the relative changes in galactosylation and afucosylation indices, two metrics that influence an antibody’s pharmacodynamics and pharmacokinetics. The N-GLYcanyzer PAT system will allow for developing a fundamental understanding of the intra/extra-cellular pathways impacting protein glycosylation dynamic flux during both fed-batch and perfusion bioprocessing. Further, such a PAT will allow the development of advanced process control strategies that can autonomously adapt to undesirable process perturbations, such as pH and temperature shifts, as well as desirable perturbations, such as the addition of specific nutrients and media modulators (e.g., sugars, cofactors), that affect the glycosylation pathway to impact drug quality.

## Author contributions

*Aron Gyorgypal*: Conceptualization, Investigation, Methodology, Validation, System Set-up, Formal Analysis, Writing – Original draft, Writing: review & editing. *Oscar Potter*: Investigation, Methodology, Conceptualization, Supervision, Writing – review & editing. *Antash Chaturvedi*: Investigation, Writing – review & editing. *David N. Powers*: Investigation, System Set-up, Validation, Writing – review & editing. *Shishir P.S. Chundawat*: Conceptualization, Investigation, Supervision, Writing – review & editing

## Acknowledgments

This work was supported by the U.S. Food and Drug Administration (FDA) through the FDA-CBER Award 1R01FD006588, Agilent Technologies University Relations Grant 4481, PC5.2-112 NIIMBL Project Award, as well as supported in part by the appointment of Aron Gyorgypal to the Research Participation Program at FDA, administered by ORAU through the U.S. Department of Energy Oak Ridge Institute for Science and Education (ORISE). The authors would like to thank Agilent Technologies Inc. (Mr. Wayne Heacock, and Dr. Ace Galermo) for their extensive and timely support of this project. As well as the Office of Biotechnology Products (OBP) Bioprocessing Lab (Dr. Cyrus Agarabi, Dr. Erica Fratz-Berilla, Mrs. Casey Kohnhorst) at the U.S. FDA Center for Drug Evaluation and Research (CDER) for their support in this project. The authors also thank GenScript Biotech Corporation (Piscataway, NJ) for the Trastuzumab cell line gift to Rutgers University.

