## Supplemental Information for "Automated Workflow for Instant Labeling and Real-Time Monitoring of Monoclonal Antibody N-Glycosylation"

**Supplementary Figures Legend**

**Supplementary Fig. 1.** Sensitivity analysis of Trastuzumab glycosylation using varying concentrations of mAb for analysis. The figure depicts fluorescence intensity as a function of concentration between a low limit of detection from mAb at 0.05 g/L upwards of 4.0 g/L. Samples were prepared from the same 4 g/L stock making dilutions with 50mM HEPES to reach each desired concentration prior to the IPC Kit chemistry sample preparation and analysis. Relative quantitation is possible at the lower concentrations using IPC chemistry.

**Supplementary Fig. 2.** FLR chromatography of VRC01 neutralizing antibody sample from offline HPLC analysis on LC-MS system prepared by the N-GLYcanyzer system. Proof of concept results show that automated N-GLYcanyzer sample preparation workflow allows for sialylated glycoforms detection. Glycoform peaks were identified based on MS analysis as described within the main text. Residence times are marginally altered as samples were run on a different LC.

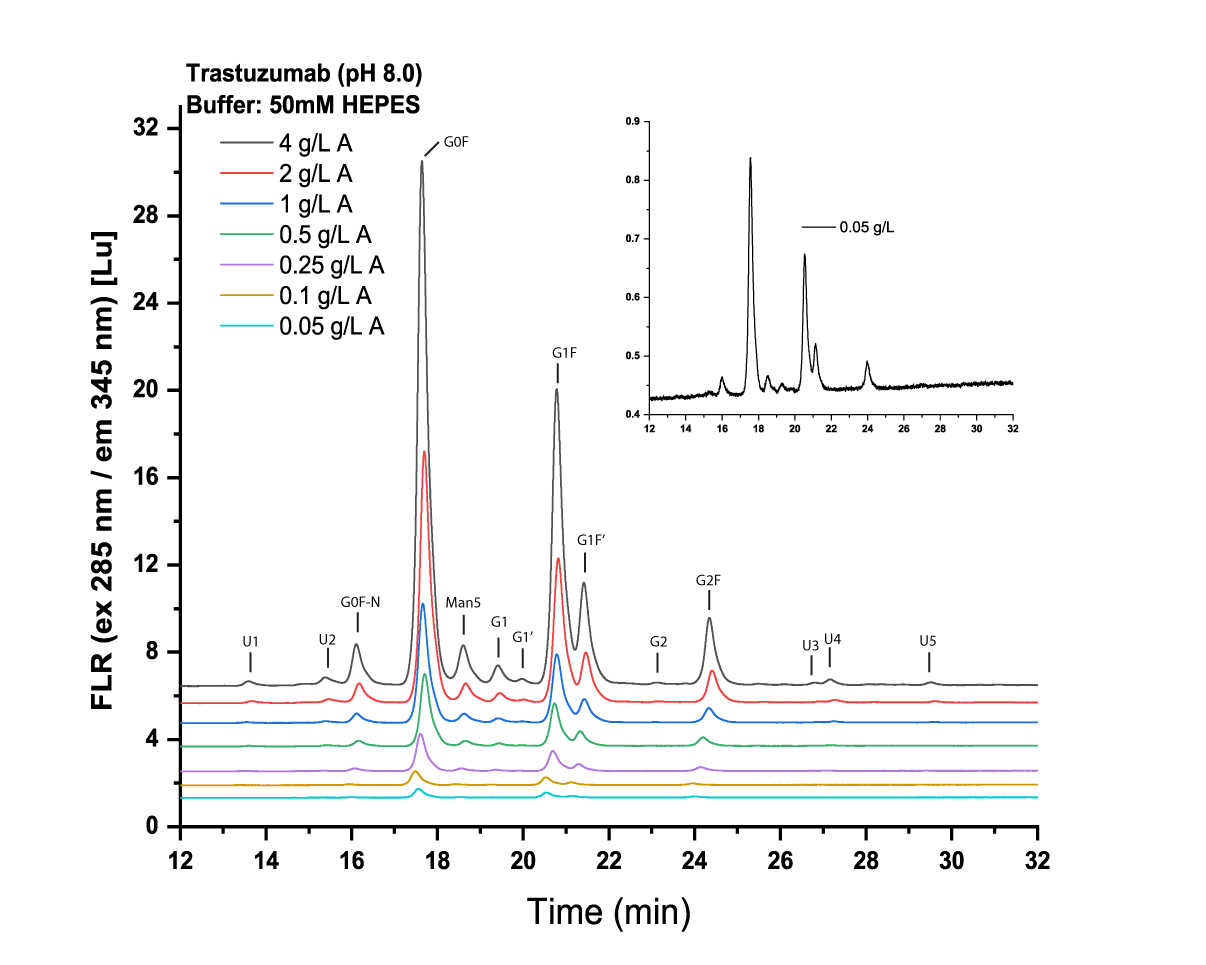

**Supplementary Fig. 1.**

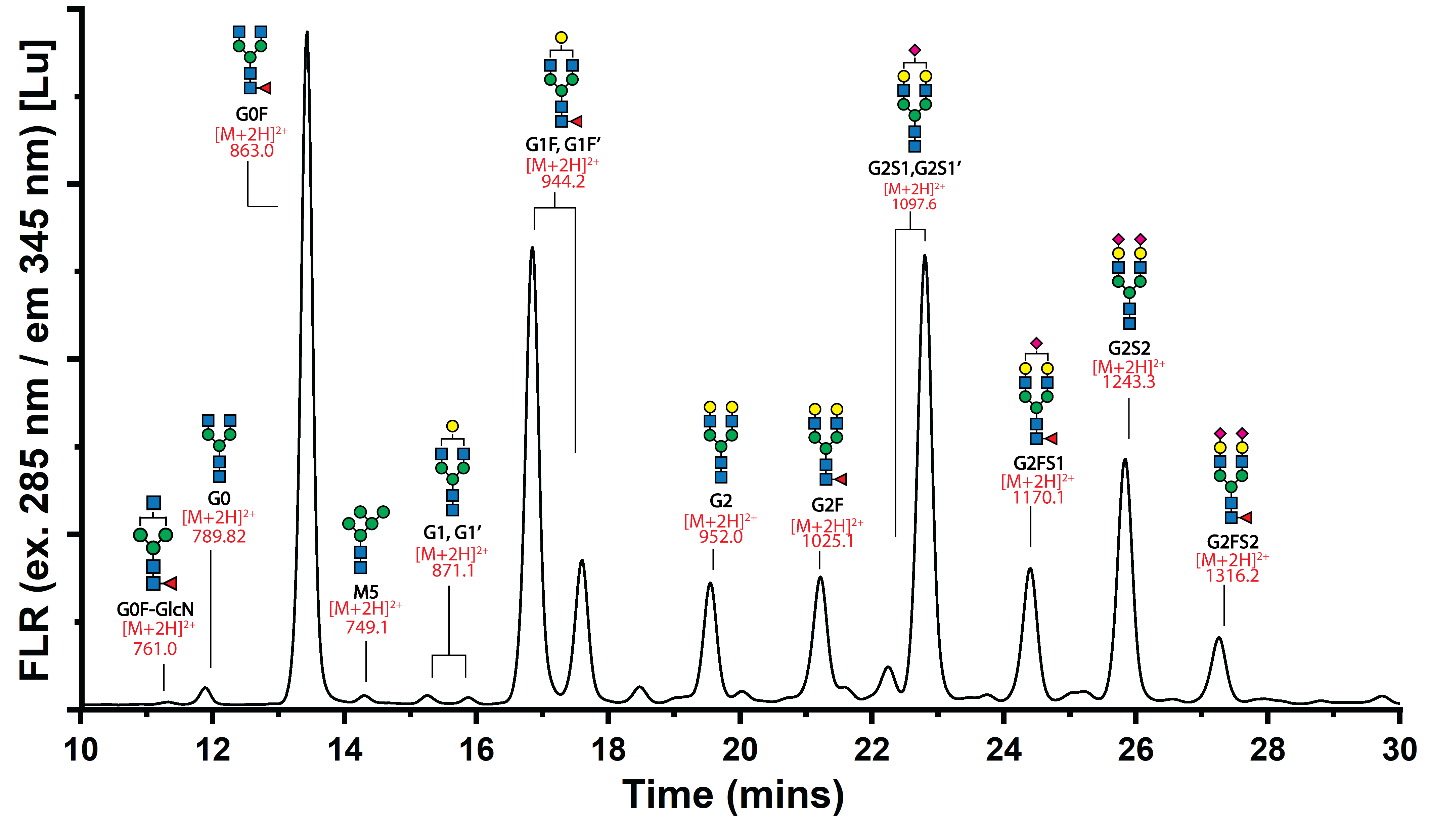

**Supplementary Fig. 2.**
